# Cellular Uptake of His-Rich Peptide Coacervates Occurs by a Macropinocytosis-Like Mechanism

**DOI:** 10.1101/2022.08.04.502757

**Authors:** Anastasia Shebanova, Quentin Perrin, Sushanth Gudlur, Yue Sun, Zhi Wei Lim, Ruoxuan Sun, Sierin Lim, Alexander Ludwig, Ali Miserez

## Abstract

Coacervates are dense microdroplets formed by liquid-liquid phase separation (LLPS) of macromolecules that have gained increasing attention as drug delivery vehicles. Recently, we have reported a new intracellular delivery system based on self-coacervating histidine (His)-rich beak peptides (HB*pep* and HB*pep*-SP) inspired by beak proteins of the Humboldt squid. These peptide microdroplets combine excellent encapsulation efficiency of therapeutics with high transfection rate and low cytotoxicity. However, the mechanism by which they cross the cell membrane remains elusive. Previous inhibitor studies provided incomplete clues into the detail uptake pathway, although they suggested a cholesterol-dependent and, possibly, an energy-independent non-classical mechanism of internalization. In this study, we improved our understanding of coacervates/cell membrane interactions using model membranes, namely Giant Unilamellar Vesicles (GUVs) and Giant Plasma Membrane Vesicles (GPMVs). We also employ a combination of electron microscopy techniques to gain detailed structural insights into the cell uptake of HB*pep* and HB*pep*-SP coacervates. We demonstrate that modulating lipid charge and cholesterol level influence coacervate attachment to GUVs. However, they are not able to cross the GUV’s lumen in an energy-independent manner. We then show that the coacervates enter HeLa and HepG2 cells via a mechanism sharing morphological features of macropinocytosis and phagocytosis, in particular involving cytoskeleton rearrangement and capture by filipodia. Our study provides key insights into the interaction of HP*pep* and HB*pep*-SP coacervates with model membranes as well as their cellular uptake pathway.

## Introduction

Coacervates, or dense liquid droplets formed by liquid-liquid phase separation (LLPS), have gained increasing attention as novel delivery agents, including for intracellular delivery^1^. Both complex coacervates (when the droplets are made by at least two oppositely charged macromolecules) or simple component coacervates are being explored^2^. Promising examples recently developed by our team are the histidine (His)-rich beak peptides – HB*pep*^3,4^ and HB*pep*-SP^5^ – derived from His-rich proteins of the Humboldt squid (*Dosidicus gigas*) beak^6^, which undergo LLPS to form simple coacervates at physiological conditions. The ability of these simple coacervate microdroplets to recruit a wide range of both low molecular weight (MW) drugs as well as large biomacromolecules, their high transfection rates and low biotoxicity hold promises to offer a real and superior alternative to traditional lipid-based delivery systems.^7^ However, mechanistic details of the cellular uptake process are still not understood. Answering this question may help to achieve better control over cellular uptake and pave the way towards more advanced peptide coacervate delivery systems and clinical translation.

Cellular uptake of sub-micron cargos is generally achieved by a variety of specialized endocytic mechanisms^8^, including clathrin mediated endocytosis (CME), clathrin-independent, fast endophilin-mediated endocytosis (FEME), clathrin-independent/dynamin-independent (CLIC/GEEC) endocytosis, caveolin-mediated endocytosis, macropinocytosis, and phagocytosis.^9^ Larger objects (> 1 μm), in contrast, have limited routes of entry into the cell. Macropinocytosis^10^ and phagocytosis^11^ are two such common endocytic routes for the entry of large particles. Phagocytosis is mostly attributed to specialized cells while macropinocytosis relates to non-specific uptake of fluidic phases and can be stimulated by various factors including growth factors.^12^ On the other hand, cell pathogens, viruses and bacteria – whose size can vary from tens of nanometres to several microns– are known to hijack cellular systems and promote their uptake after engaging with the cell surface. Examples may include macropinocytosis-like mechanism or surface mimicry and hijacking of receptor-mediated endocytosis.^13,14^ Such non-canonical pathways often share some features with common entry routes but can be distinctly regulated.^15^

Although there are recent reports of coacervates designed for drug or gene delivery applications, in most cases their uptake mechanism has not been investigated in detail. Iwata *et al.*^16^ proposed a macropinocytosis-like mechanism for the uptake of complex micron-size peptide-based droplets, mostly based on inhibitor studies indicating involvement of cytoskeleton rearrangement. The non-selectivity of macropinocytosis could partially explain why micron-sized coacervates could be internalized via this route since most coacervate delivery systems reported in the literature lack specific ligands required for interacting with the cell surface. Other examples of coacervates found to be internalized by cells include those based on polycation-b-polypropylene glycol di-block copolymer complexed with heparin,^17^ as well as dextran-based^18^ or starch derivatives^19^ nanocoacervates. However, their uptake mechanism has not been addressed. Armstrong *et al.* ^20^ reported the selective delivery of complex coacervates composed of ATP and cationic poly- (diallyldimethylammonium chloride) (PDDA) via direct microdroplet-cell fusion. Since there is evidence of both energy-dependent and energy-independent uptake, it appears that the internalization route of coacervates may depend on the chemical nature of the coacervates themselves. It should also be noted that all of the above reports involve complex coacervates, for which charge-charge interactions can lead to full internalization into liposomes under certain conditions.^21^ In contrast, the internalization into GUV’s and the cellular uptake of single-component peptide or protein coacervates remains unknown. In addition to electrostatic interactions, other type of interactions may be involved in their attachment to the cell membrane. In particular, aromatic residues that are often enriched in simple coacervates tend to be present in the interfacial region of lipid membranes^22^ and are usually involved in protein/lipid bilayer interactions ^23^ and in peptide/protein membrane anchoring^24^. For example, tuning the properties of aromatic rings has been shown to increase the cellular uptake of peptides^25^.

HB*pep* and HB*pep*-SP peptide are simple coacervates that are enriched not only in His but also in aromatic residues.^5^ Based on results from pharmacological inhibitors, where only MβCD and low (4°C) temperature prevented coacervate delivery into cells whereas sodium azide (NaN3) and other endocytic inhibitors did not affect uptake^5^, we concluded that the uptake could occur via non-classical endocytosis and may involve cholesterol-rich lipid rafts on the membrane. We cautiously speculated that the coacervates could interact and directly fuse to the plasma membrane at these cholesterol-rich regions, not unlike the coacervate-cell fusion reported by Armstrong *et al.*^20^ However, the details of the uptake mechanism of HB*pep* and HB*pep*-SP remained unclear. Notably, a chemical inhibitor analysis of cellular uptake of polystyrene-co-poly(N-) isopropylacrylamide microgels with different stiffness provided evidence of phagocytosis for macrophages, but did not affect uptake in the presence of all inhibitors in hepatocarcinoma cells. The uptake was only affected by cold temperature.^26^ It might be that the internalization of micron-sized objects by non-phagocytic cells, like in case of pathogens, shares some features of common uptake routes but is biochemically distinct. Alternatively, the common protocols for inhibitor assays are not applicable in some cases.

In this report, we show that HB*pep* and HB*pep*-SP coacervates are in fact taken up by a mechanism that shares some morphological features of macropinocytosis and phagocytosis. Studies with simple model membrane systems such as Giant Unilamellar Vesicles (GUVs) and Giant Plasma Membrane Vesicles (GPMVs) revealed that coacervates merely attached to the GUV and GPMV surfaces but failed to cross into the lumen, suggesting an energy-dependent and possibly a cytoskeletal-requiring uptake mechanism. Further experiments exploiting direct visualization of cellular uptake of coacervates by transmission and scanning electron microscopy (TEM and SEM) allowed to unambiguously visualize the membrane engulfment and endocytosis of coacervates in two different non-phagocytic cell lines, *i.e.* HeLa and HepG2. We were able to capture different stages of HB*pep* and HB*pep*-SP coacervates internalization, from attachment to membrane wrapping to complete membrane engulfment. At the attachment stage, involvement of cell membrane’s filopodia in capturing the coacervates was observed, which is likely a critical step towards their subsequent internalization. Our study provides new understanding into the interaction of His- and aromatic-rich peptide coacervate microdroplets with model membranes and their subsequent cell uptake via non-classical endocytosis.

## RESULTS AND DISCUSSION

### Interactions between HB*pep* coacervates and lipid membranes with GUVs and GPMVs

#### Charge-charge interactions

Cellular uptake of most nano- or microparticles begins with their adhesion to the outer leaflet of the plasma membrane and is often dictated by electrostatic interactions. To determine whether charge-charge interactions played a role in the initial step of coacervate adhesion to lipid membrane, HB*pep* coacervates entrapping enhanced Green Fluorescence Proteins (EGFP) were mixed with GUVs prepared from the zwitterionic phospholipid 1-palmitoyl-2-oleoyl-glycero-3-phosphocholine (POPC) (**Figure 1A,** chemical structure). GUVs are simplified model membrane systems with tunable physio-chemical properties that make them a convenient tool for studying membrane-coacervate interactions^27^. HB*pep* coacervates when mixed with POPC GUVs at three different pHs (7.5, 8.5 and 9.5) exhibited a pH-dependent effect in its surface attachment to GUVs (**Figure 1A,** bar graph). The highest level of coacervate attachment was observed at pH 7.5 with 0.54 (± 0.31) coacervates/μm of GUV, but this attachment decreased to 0.13 (± 0.04) and 0.16 (± 0.1) coacervates/μm of GUV at pH 8.5 and 9.5, respectively.

**Figure 1.**
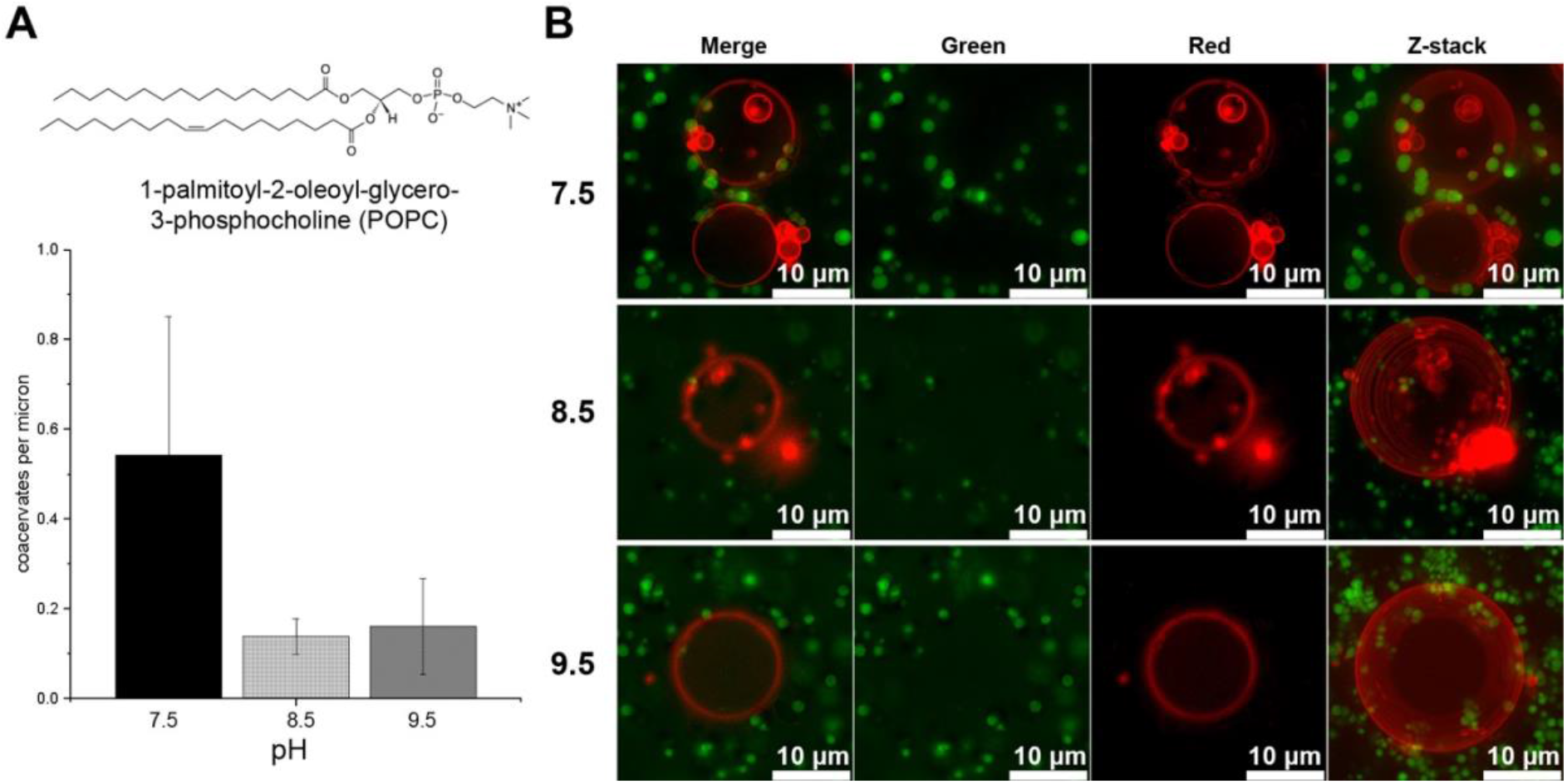
Effect of pH on HB*pep* coacervate attachment to POPC GUVs. **(A)** Plot of coacervates attached per micrometer of GUV, indicating highest coacervate attachment at pH 7.5. **(B)** Fluorescence microscopy images of POPC GUVs (Red) mixed with HB*pep* coacervates entrapping EGFP (green) at different pHs. Representative merged images of the green and red channel (middle columns) are shown in the left-most column and the right-most column represents a representative Z-stack composite of optical sections collected 1 μm apart.

The decreased attachment at higher pHs could partly be due to the peptide acquiring a net negative charge that in turn affects the surface properties of the coacervates, as seen by a change in the size and morphology of coacervates (**Figure 1B**, green channel). HB*pep* peptide is zwitterionic with charge contributions from five His residues in its sequence and two oppositely charged termini, resulting in a theoretical pI of 7.97. Thus, they are partially positively charged at pH 7.5, which we expect slightly facilitate coacervate attachment to the outer membrane of the GUV and by extension, to the negative charges in the outer leaflet of the plasma membrane.

Further exploration of the role of electrostatic interactions at the GUV membrane surface was obtained by varying the ionic strength of the medium when HB*pep* coacervates were mixed with POPC GUVs at pH 7.5. HB*pep* coacervates exhibited relatively poor attachment to GUVs at low (≤ 58 mM) salt concentrations (data not shown). An initial increase in salt concentration to 0.1 or 0.15 M led to a corresponding increase (0.54 [± 0.30] and 0.52 [± 0.12] coacervates/μm of GUV, respectively) in coacervate attachment (**Figure 2A and Suppl. Figure. S1**). However, at higher ionic strengths (≥ 0.2 M), the trend was reversed and had a negative effect on coacervates attachment (0.20 [± 0.10] and 0.14 [± 0.09] coacervates/μm of GUV) (**Figure 2A**).

**Figure 2.**
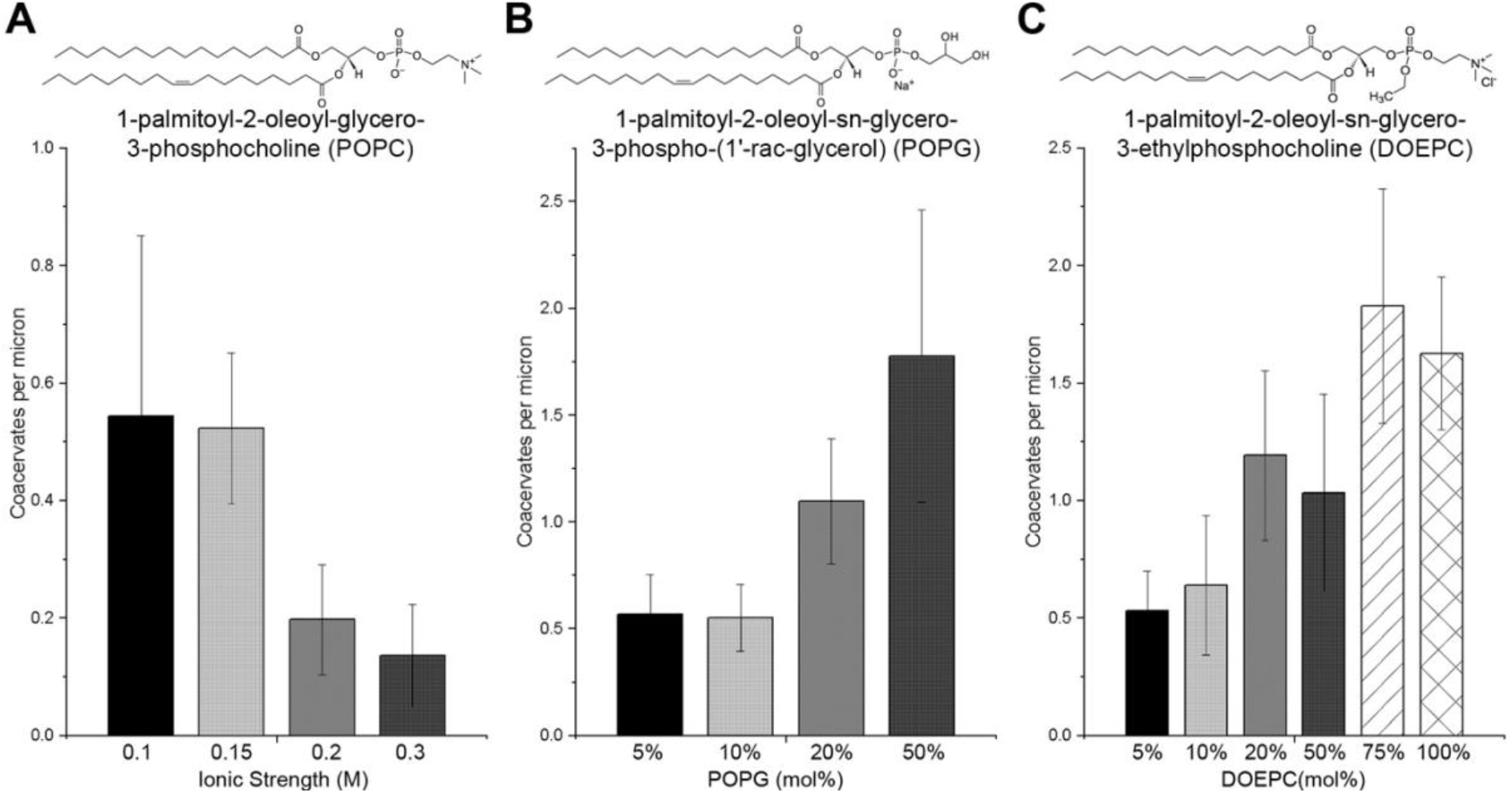
Effect of ionic strength and lipid charge on HB*pep* coacervate attachment. Plot of coacervate attachment to **(A)** POPC GUVS prepared at different ionic strengths showing less attachment as ionic strength increases, **(B)** POPC GUVs with increasing amounts of negatively charged POPG and **(C)** positively charged DOEPC showing that coacervate attachment increases when the levels of charged lipids increases in GUVs.

Finally, mixing HB*pep* coacervates with POPC GUVs containing increasing amounts of positively or negatively charged phospholipids showed that HB*pep* coacervates attached more to GUVs composed of charged lipids than to zwitterionic lipids (**Figure 2B-C and Suppl. Figures S2&3**). In general, coacervate attachment increased with increasing amount of 1-Palmitoyl-2-oleoyl-sn-glycero-3-phospho-rac-(1-glycerol) (POPG), attaining a maximum of 1.78 (± 0.69) coacervates/μm of GUV when the POPG concentration was 50 mol% (**Figure 2B**). Similarly, coacervate attachment increased with increasing amount of 1,2-dioleoyl-sn-glycero-3-ethylphosphocholine (DOEPC), reaching a maximum of 1.83 (± 0.50) coacervates/μm of GUV when the DOEPC concentration was 75 mol% (**Figure 2C**), indicating that coacervates preferred adhering to membranes with charged lipids regardless of the type of charge. Although electrostatic interactions are obviously of great importance, aromatic-rich peptides add complexity to cell/peptide coacervate interactions. Andreev *et al*.^28^ showed that hydrophobic interactions may govern the adhesion of aromatic-rich peptoids to charged bilayers via a two-step mechanism, *i.e.: (i)* transfer from to the aqueous environment to interfacial regions, and (*ii)* further penetration into the lipid core. While peptoids with 30-50% aromatic content only caused deep lipid penetration of charged membranes and interacted with zwitterionic/cholesterol membranes in the interfacial region, further increase in aromatic content led to the penetration and disrupting of both types of model membranes. Therefore, in the case of HB*pep* coacervates their stronger preference to charged membranes might be related to their elevated aromatic content (23 mol% of aromatic amino acids).

The above set of results indicate that under physiological conditions, HB*pep* coacervates adhere to 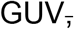 but none of the HB*pep* coacervates were able to enter the GUV lumen, suggesting that adhesion energy alone may be insufficient for HB*pep* coacervates to bend the lipid membrane till the full engulfment occurs, and that there may be additional factors or conditions required for completing the uptake process.

#### Membrane fluidity

Cholesterol (**Figure 3A**) is an integral component of eukaryotic cell membranes and is a key molecule in controlling membrane fluidity, *i.e.* membranes with a high cholesterol content exhibit decreased membrane fluidity. Cholesterol concentration within the cell membranes can range anywhere from < 5 mol% in mitochondrial membranes up to 25 mol% in plasma membranes.^29^ To evaluate the role of membrane fluidity in coacervate attachment, HB*pep* coacervates were mixed with POPC GUVs prepared with cholesterol whose concentration was varied between 10 to 50 mol%. Interestingly, HB*pep* coacervates exhibited an appreciable increase in attachment only for POPC GUVs containing 20 mol% cholesterol (0.75 [± 0.19] coacervates/μm of GUV, **Figure 3B-C**), whereas POPC GUVs containing 10, 30, 40 and 50 mol% cholesterol resulted in decreased coacervate attachment (**Figure 3B**). At higher cholesterol levels (above 30%), coacervates attachment decreased to below 0.40 coacervates/μm, suggesting that excessive membrane fluidity or excessive membrane rigidity was not favorable for coacervate attachment. These results are in agreement with inhibitor studies with MβCD reported in our previous work^5^, where coacervate uptake was abolished in the presence of MβCD suggesting a possible role for lipid raft mediated uptake. However, additional experiments with lipids rafts that were spontaneously induced on GUVs using a 40% POPC/40% SM/20% cholesterol lipid composition did not result in coacervate uptake into GUVs **(Supplementary figure S5)**. It is possible that a specific range of fluidity modulated by cholesterol is favorable to anchor the aromatic amino acids to the cell membrane, and thus facilitates coacervate attachment which, in turn, promotes their uptake.

**Figure 3.**
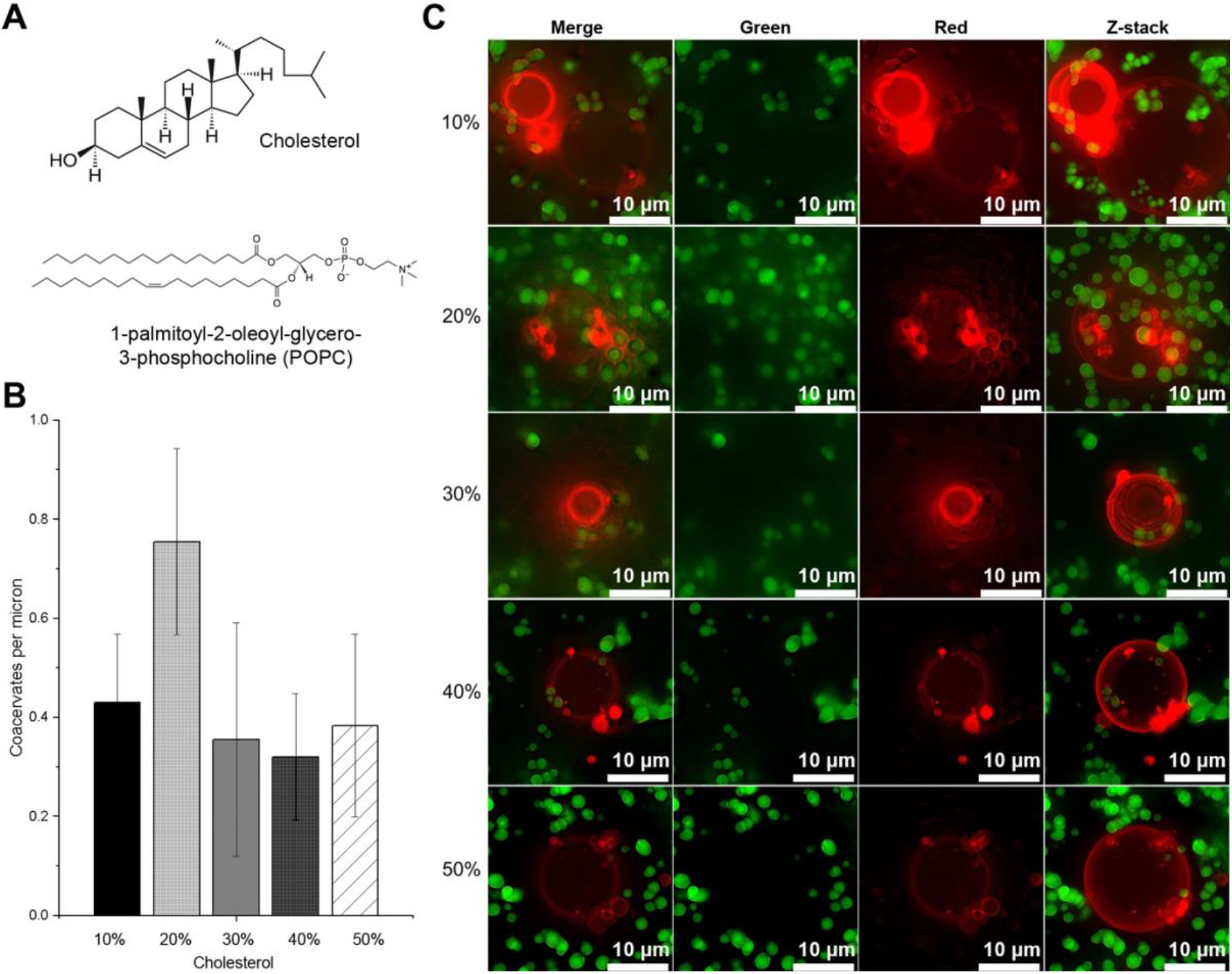
Effect of membrane fluidity on HB*pep* coacervate attachment. **(A)** Chemical structures of cholesterol and POPC employed in preparing the GUVs. **(B)** Plot of coacervates attached per micrometer of GUV, indicating that coacervate attachment is highest when POPC GUVs were prepared with 20% cholesterol. **(C)** Fluorescence microscopy images of POPC GUVs with varying cholesterol content (Red) mixed with HB*pep* coacervates entrapping EGFP (green). Representative merged images of the green and red channel (middle columns) are shown in the left-most column and the right-most column represents a representative Z-stack composite of optical sections collected 1 μm apart.

While the above results indicate that a certain cholesterol content is preferred for coacervates attachment, the latter did not cross the membrane into the GUV lumen no matter the cholesterol concentration. Albeit robust membrane model systems, GUVs do not represent the full complexity of actual cellular membranes. Thus, we conducted further studies by substituting GUVs with GPMVs, as described below.

GPMVs offer a few advantages over GUVs in terms of having a physiological lipid composition containing native proteins including glycosyl-phosphatidyl-inositol-anchored proteins. They also have a smaller order difference between lipid phases, contain cholesterol, and may preserve some level of membrane asymmetry.^30^ However, similar to GUVs, they lack a cytoskeletal support that can be used to distinguish the contribution of the plasma membrane from that of active processes. GPMVs prepared from HeLa cells (**Figures 4A-B**) when mixed with HB*pep* coacervates loaded with EGFP yielded similar results to those seen with GUVs, namely coacervates attached to GPMV surfaces but were not internalized, suggesting that the uptake process may be energy-dependent and/or requires a cytoskeletal system (**Figure 4C-E**). While cell uptake assessment with inhibitors may provide clues on the uptake mechanism, the inhibition is often not very specific and can be cell type dependent.^31^ Moreover, blocking of one endocytic pathway can lead to the upregulation of others.^32^ To avoid such pitfalls, we decided to investigate the uptake process in cells using TEM and SEM.

**Figure 4.**
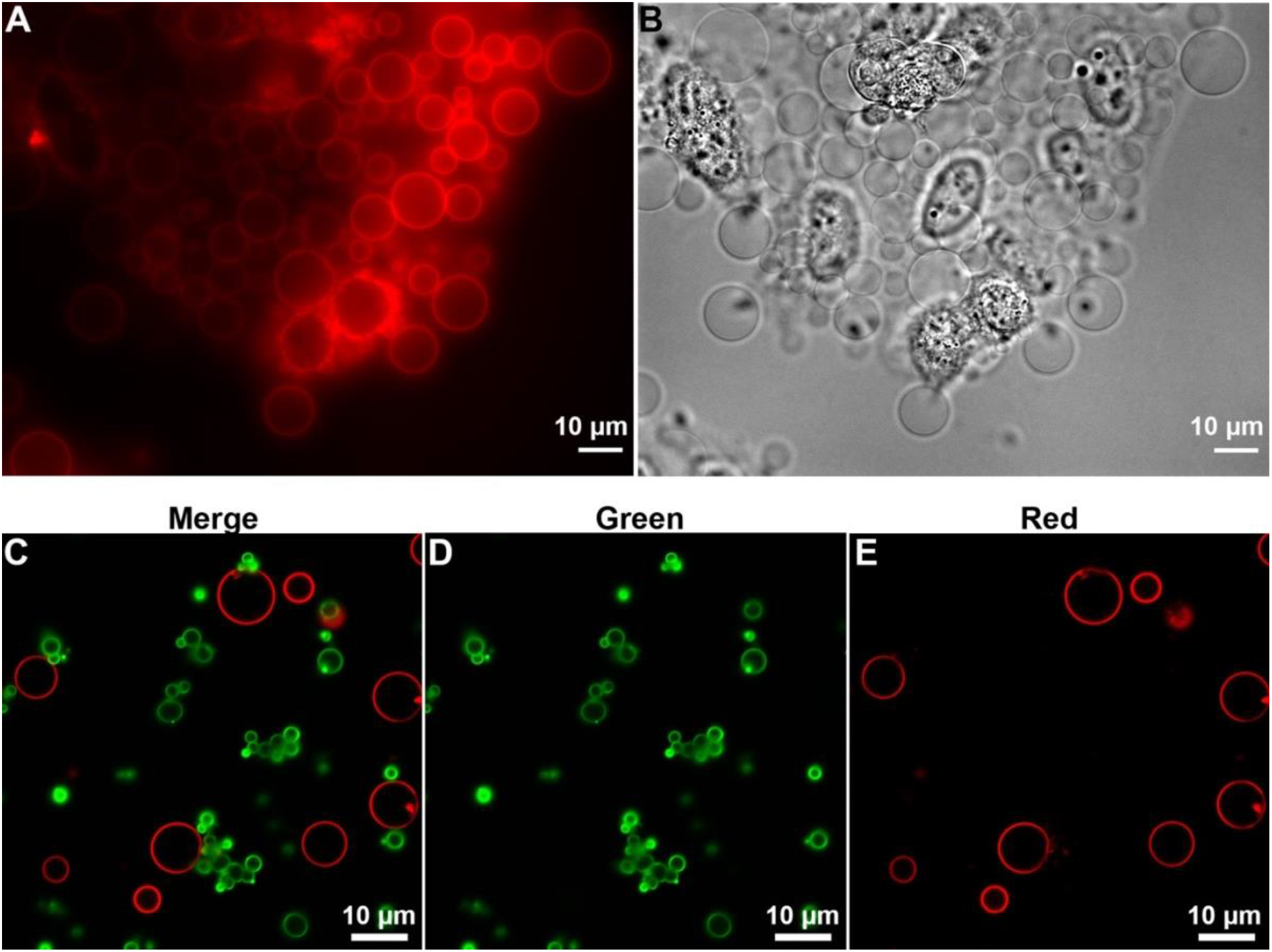
HB*pep* coacervate interaction with GPMVs. **(A)** Fluorescence and **(B)** brightfield microscopy images of GPMVs (red) prepared from HeLa cells. **(C)** confocal images of HB*pep* coacervates trapping EGFP (green) mixed with GPMVs (red) show coacervate attachment to GPMVs but no uptake. **D & E** are images captured in the green and red channel, respectively.

### Cellular uptake of HB*pep* and HB*pep*-SP coacervates in HeLa and HepG2 cells: TEM studies

Although HB*pep* and HB*pep*-SP coacervates are micron-size objects, making them in principle suitable to be observed by fluorescence microscopy, we found that they were able to recruit many dyes (including membrane dyes, data not shown). As a result, imaging of coacervate-cell membrane interactions by fluorescence microscopy proved challenging with ambiguous outcomes. Therefore, we relied on a combination of TEM and SEM imaging techniques as an alternative. Hence, ultrathin (70-100 nm) sections of fixed and resin embedded HeLa cells treated with HB*pep* coacervates for 3 h were imaged using the High-Angle Annular Dark-Field Scanning TEM (HAADF-STEM) technique. **Figure 5** is a panel of HAADF-STEM images that shows micron-sized spherical droplets, representing HB*pep* coacervates, at various stages of the uptake process: (*i)* outside the cell, (*ii*) partial membrane engulfment, (*iii*) an advanced stage of cup-shaped extensions whose leading rims appear to close over the coacervates, and (iv) internalized coacervate in the cell cytoplasm. Since HAADF-STEM is a dark-field technique where the objects containing heavy elements appear brighter on the image, cell membranes stained with osmium tetroxide (OsO4) appear bright on the dark background. The high electron density observed for the coacervates is likely due to the high peptide concentration within the coacervates as well as heavy metal enrichment during staining (**Figures 5 A-D**). Such spherical droplets were not observed in control cells **(Supplementary figure S5)**. Besides being electron dense, some coacervates were slightly elongated **(Figure 5C)** which can be attributed to their deformability. Additionally, fully membrane-enclosed coacervates in the form of endocytic vesicles were seen within the cytoplasm (**Figures 5 A,D**). Hence, there is evidence that HB*pep* coacervates are taken up by endocytosis. The cup-shaped structure of the plasma membrane identified in **Figures 5 B-C** is a characteristic feature of macropinocytosis and phagocytosis^12^ and involves actin polymerization and reorganization. In general, the process of macropinocytosis is morphologically very similar to that of phagocytosis. However, in the latter case, receptors on the plasma membrane engage with solid particles (that are about to be engulfed) to guide the cup formation^33^. Macropinocytosis, instead, relies on flat membrane protrusions that occasionally fold back to form circular ruffles. The resulting macropinosomes usually contain some amount of liquid surrounding internalized particles, whereas during phagocytosis a tight sleeve is formed around internalized object. In the case of HB*pep* coacervates, we observed a tight connection between the membrane and the surface of internalized coacervates, which is more typical for phagocytosis (**Figure 5D**). A recent study utilized coacervates as a template for lipid bilayer self-assembly^34^ and demonstrated a high affinity for lipids towards coacervates. In the case of HB*pep* coacervates, strong adhesion to the lipid bilayer could be partially related to their enrichment in aromatic acids. Notably, formation of “lipid corona” was identified for gold nanoparticles decorated with aromatic amino acids interacting with lipid vesicles^35^.

**Figure 5.**
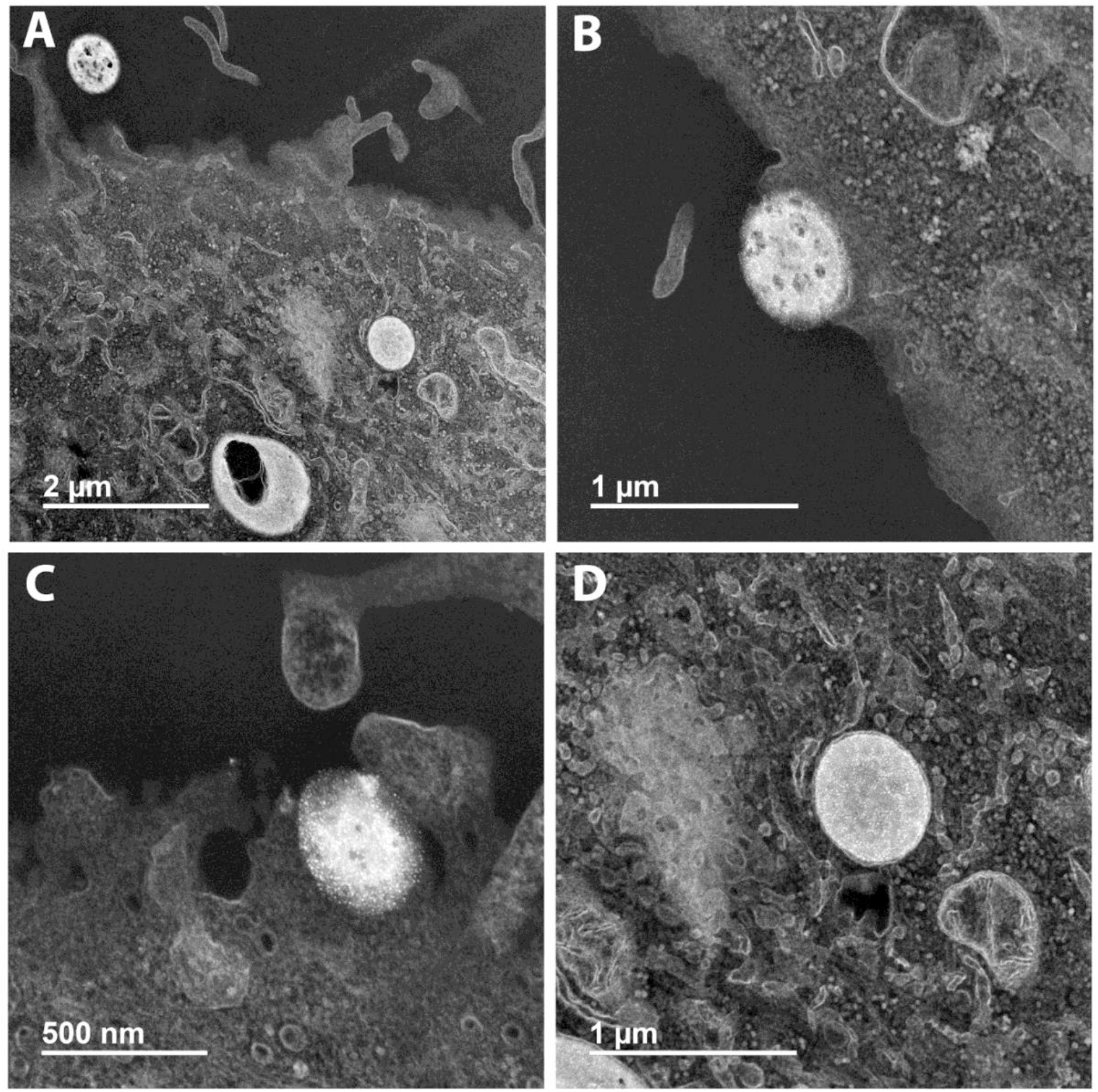
HAADF-STEM images of HB*pep* coacervates in HeLa cells. Ultrathin sections of fixed and resin embedded HeLa cells showing HB*pep* coacervates during the various stages of cell uptake. **(A)** coacervate droplets outside the cell and already internalized and presented in cell cytoplasm; **(B,C)** different stages of membrane engulfment around the droplet; **(D)** fully internalized coacervate in the cell cytoplasm.

We further extended our studies to HB*pep*-SP peptide coacervates –in which one extra Lys residue is modified with a disulfide-containing self-immolative moiety (Table 1)– to two cell lines, namely HeLa and HepG2 **(Figure 6).** Unlike HB*pep* coacervates, the HB*pep*-SP peptide coacervates undergo disassembly once inside the cells triggered by reduction of the disulfide bond located in the moiety attached to the Lys residue, with concomitant release of payload.^5^

**Figure 6.**
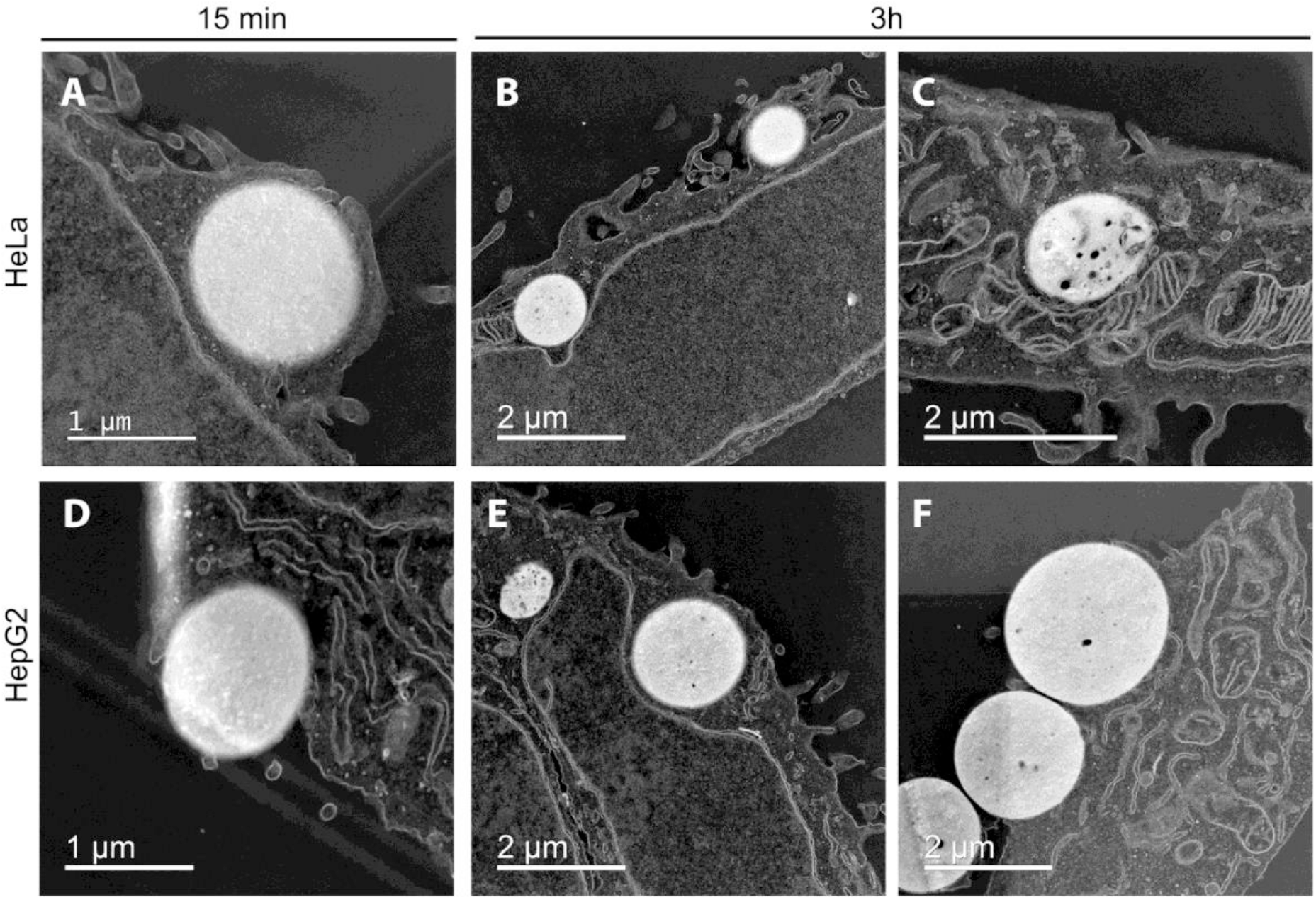
HAADF-STEM images of HB*pep*-SP coacervates in HeLa and HepG2 cells. Ultrathin sections of fixed and resin embedded HeLa (**A-C**) and HepG2 (**D-F**) cells showing HB*pep-*SP coacervates during the various stages of endocytosis after 15 min (A,D) and 3h (B, C, E, F) of incubation time. Cup-like structure (A) and sinking behaviour (D,F) is observed. After 3h of incubation, not all of the coacervates were fully internalized.

**Table 1.**
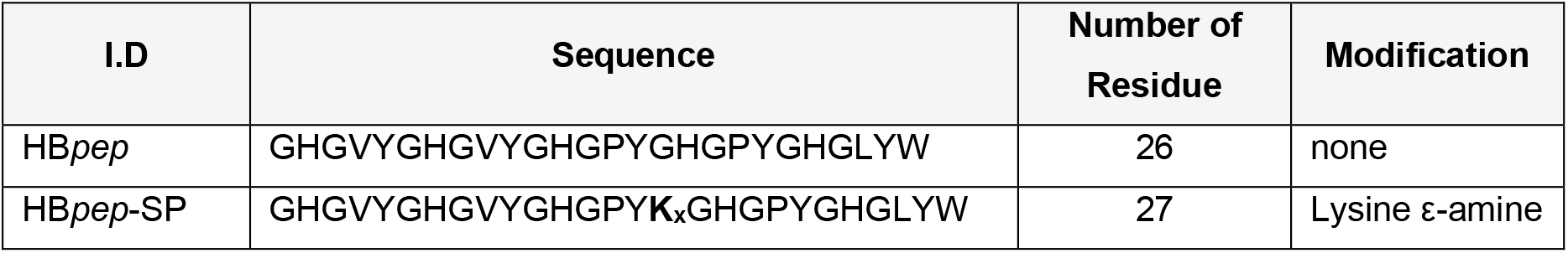
Amino acid sequence of HB*pep* and HB*pep*-SP

Similar to HB*pep* coacervates, multiple electron dense coacervates were observed, demonstrating membrane engulfment and endocytosis in both cell lines (**Figure 6**). Micrometer-sized droplets attached to the cell membrane as well as fully internalized droplets within the cells could be seen after 15 min **(Figure 6 A,D)** and 3 h **(Figure 6 B,C,E,F)** of incubation. Since HB*pep*-SP coacervates disassemble once in the cell, we observed some signs of their dissolution (**Figure 6C**). Possibly, some of the droplets can bind to the cell surface and undergo internalization later. Notably, some of the droplets looked like they were “sinking” into the cell, bending the cell membrane with almost no pronounced membrane extensions (**Figure 6F**). Some thin filaments around the coacervates could be observed, which probably represent filopodia that are thin actin projections protruding from the cell surface to actively search for chemical and structural cues and help the cell sense its external environment.^36^ We did not notice this kind of coacervate interactions with cell membrane for HB*pep* and HeLa cells (**Figure 5**), probably because the sections were prepared from cell monolayer sectioned parallel to the substrate surface. In contrast, HB*pep*-SP cell samples were prepared as a pellet, so images of multiple cell orientations from the section could be captured.

### Characterizing spatial relationship between HB*pep* and HB*pep*-SP coacervates and plasma membrane: SEM studies

Since TEM was performed on ultrathin sections, which is not suitable to provide information about the cell surface topography, we opted for SEM to gain further insights into the internalization process and visualize the entire cup-shaped extension and engulfment of coacervates. **Figure 7** is a panel of SEM images that show spherical droplets of HB*pep*-SP on the surface of HepG2 cells after 15 min of incubation, similar to those observed with TEM **(Figure 6).** Multiple coacervates were observed interacting with the cell surface (**Figure 7A**). Some were attached to the cell and surrounded with filopodia (**Figure 7B**). We also could visualize flat membrane extensions wrapping coacervates (**Figure 7C**) as well as partial membrane engulfment of coacervates (**Figure 7D**). Similar to TEM, closer inspection of SEM images for both HB*pep* and HB*pep*-SP in HeLa and HepG2 cell lines revealed different stages of membrane engulfment (**Figure 8**). We classified the observed types of coacervate/cell interaction topologies as “adhesion”,” sinking’’ and “ruffle/cup’’. We confirmed that many coacervates were often wrapped by filopodia (**Figure 8 A-C,F-G,J**), providing an intriguing parallel with filopodia capture by some pathogens during the initial stages of attachment to the cell^15^. Ingle *et al.* (2020)^37^ showed that filopodia facilitate the internalization process of nanocomplexes by sensing and steering them into the main body of the cell, thus serving as guides for the direct endocytosis of polyplexes. It is possible that the filopodia help capture the coacervates and facilitate their internalization. On the other hand, unlike TEM, SEM does not allow to visualize fully internalized coacervates, although we could see some bumps on the cell surface (**Supplementary figure S6**), that may represent fully engulfed coacervates.

**Figure 7.**
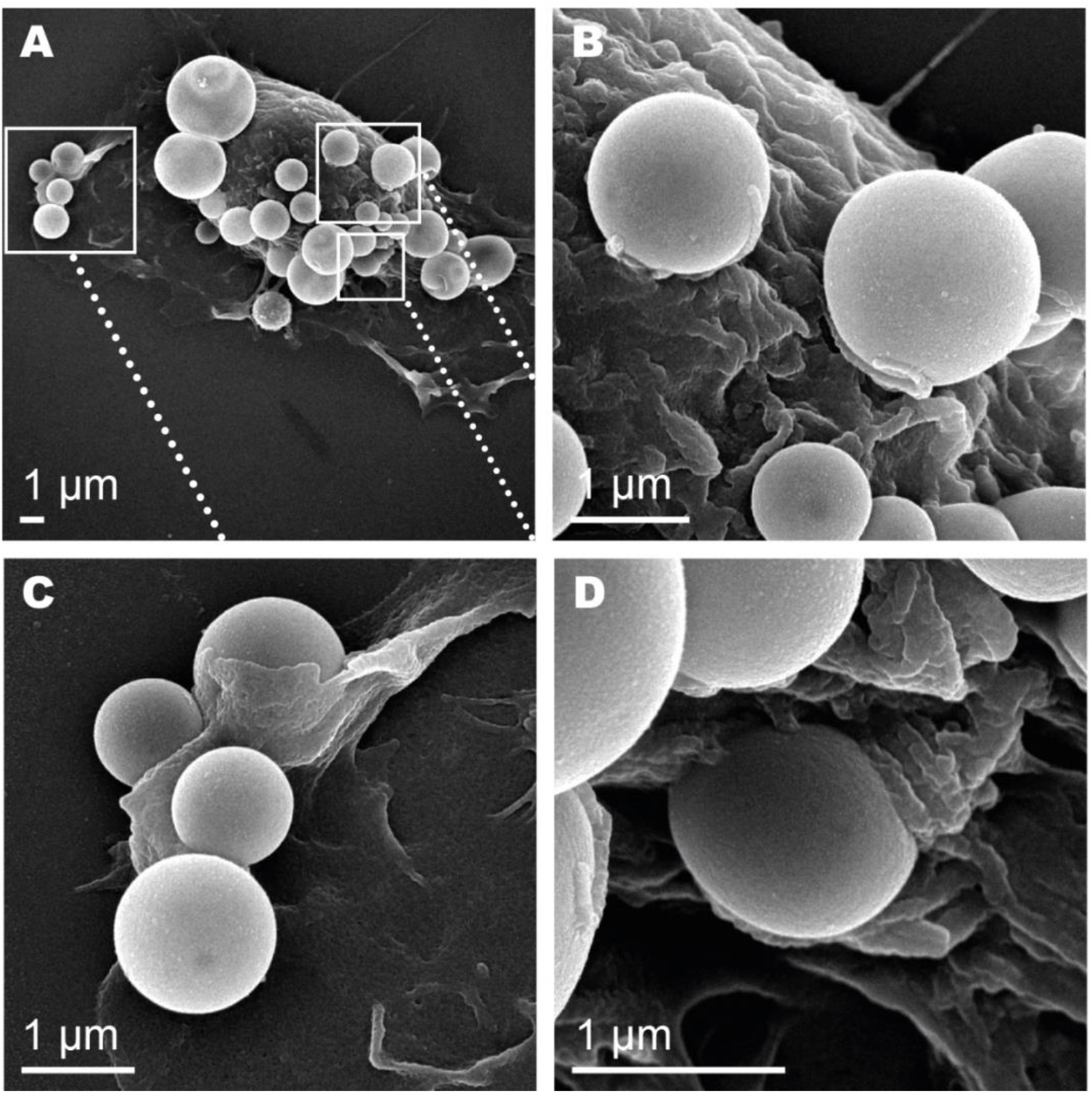
SEM images of HB*pep*-SP coacervates interacting HepG2 cell surface demonstrating various stages of uptake. **(A)** Representative SEM micrograph of a cell with multiple coacervates after 15 min of incubation. **(B)** Coacervates attached to the cell surface. **(C)** Wrapping of coacervates by flat membrane protrusion. **(D)** Partial membrane engulfment.

**Figure 8.**
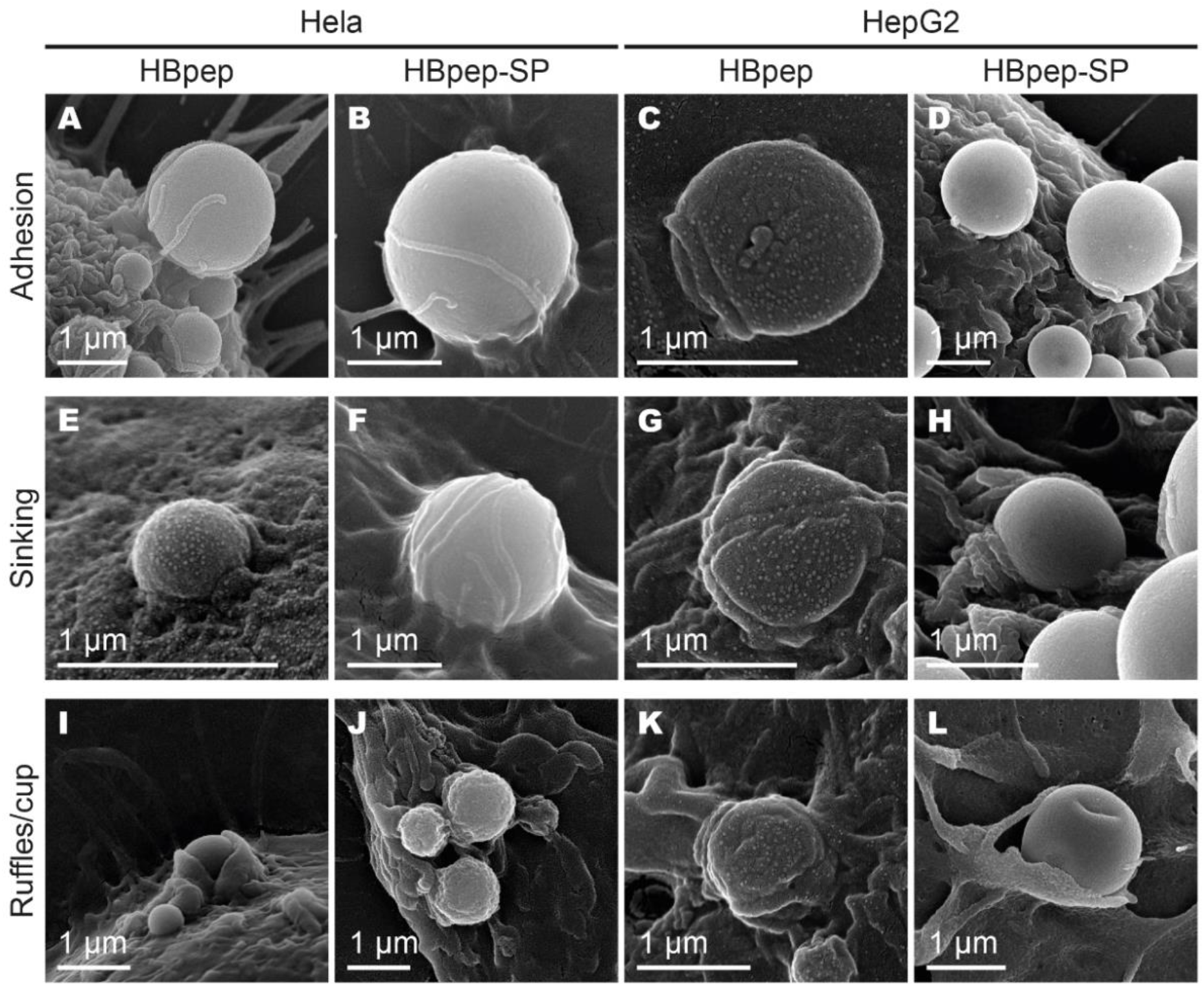
Close-up views of HB*pep* and HB*pep*-SP coacervates interacting with HeLa and HepG2 cell surface visualized by SEM. **(A-D)** adhesion; **(E-H)** sinking; (**I-L)** wrapping by flat membrane protrusion/cup-like structure.

**Figure 9** provides a comparison between TEM and SEM images of observed coacervate/cell interaction topologies on the surface of the cell: adhesion involving filopodia capture, cup-like structure and sinking of coacervate. These three steps could represent different stages of one internalization pathway or a combination of different internalization mechanisms sharing features of both macropinocytosis and phagocytosis.

**Figure 9.**
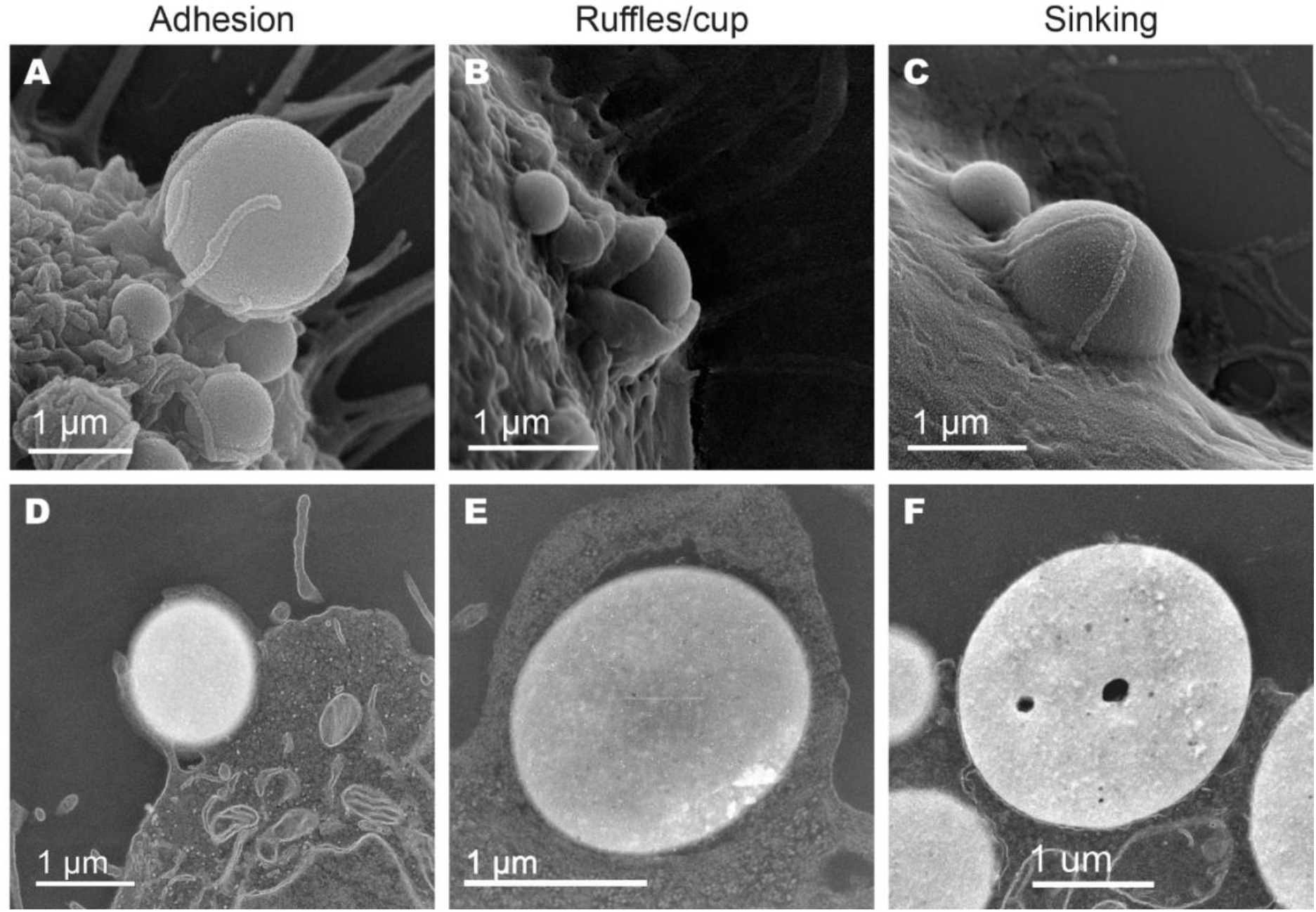
Comparison of representative (A-C) SEM and (D-F) HAADF-STEM images of HB*pep* and HB*pep*-SP coacervates interacting with HeLa and HepG2 cell surfaces illustrating the three main types of coacervates/cell interaction topologies. Adhesion and filopodia capture (A,D); ruffle/cup (B,E); sinking (C,F).

While TEM sections do not allow to clearly differentiate filopodia from membrane ruffles (although filopodia appear to be thinner) (**Figure 9 A,D**), SEM imaging enables to easily distinguish finger-like extensions of filipodia^38^ from flat membrane protrusions (**Figure 9 B,E**). An interesting example is the sinking type of behavior: while SEM visualizes the upper part of coacervates on the cell surface, TEM imaging of ultrathin sections allows to directly visualize the cell membrane, its contact with coacervates’ surfaces and membrane bending (**Figure 9F).**

## CONCLUSIONS

Using a combination of tools, we have demonstrated that HB*pep* and HB*pep*-SP coacervates are taken up by HeLa and HepG2 cell by a mechanism sharing features of phagocytosis and macropinocytosis. The data from GUVs and GPMV studies reveal adhesion of coacervates at the membranes, but no internalization. Experiments with GUVs where the pH, ionic strength of the media, amount of negatively or positively charged lipids in the GUVs and cholesterol levels were varied, indicated that a combination of electrostatic interactions with other interactions involving aromatic residues might govern the coacervates/GUVs membrane attachment. The inability of coacervates to cross the lipid bilayer and reach GUVs and GPMVs lumen suggests an energy-dependent mechanism of cell uptake.

Direct visualization of the coacervate-cell membrane interaction by HAADF-STEM and SEM allowed us to observe various stages of cellular uptake. From attachment –often accompanied by filopodia capture– to full membrane engulfment. These results suggest the active involvement of the cell cytoskeleton in coacervate internalization and lead us to revisit the previous assumptions of HB*pep* and HB*pep*-SP coacervates uptake, pointing towards an energy-dependent mechanism akin to macropinocytosis. Moreover, this study shows that the combination of HAADF-STEM and SEM techniques is a powerful tool to study the interaction of peptide coacervates with the cell surface and their subsequent internalization, providing new insights into these processes.

## MATERIALS AND METHODS

### Materials

#### Peptides

HB*pep* was purchased from GL Biochem (Shanghai) Ltd, China and subjected to an additional purification step by High Performance Liquid Chromatography (HPLC) using a C8 column. HB*pep*-SP was synthesised according to the protocol described previously.^5^ To prepare the coacervates, 10 mg/mL of HB*pep* or HB*pep*-SP in 10 mM acetic acid was pipetted into phosphate buffer (pH 6.5, 100 mM ionic strength) at a volume ratio of 1:9 as previously described.^5^

#### EGFP-loaded coacervates for GUV and GPMV’s studies

1 μL of EGFP stock solution (1 mg/mL) was added to 89 μL of 0.1 M phosphate buffer at pH 7.5 (or 0.1 M phosphate buffer at pH 8.0 for GPMV experiments). 10 μL of 10 mg/mL HB*pep* stock solution in 10 mM acetic acid was added to initiate recruitment and coacervation process.

#### Giant unilamellar vesicles (GUVs)

GUVs were prepared via gel-assisted formation on PVA. Briefly, 5% (w/w) solution of Polyvinyl alcohol (PVA) was prepared by stirring PVA in water at 90 ^o^C. 100 μL of PVA solution was added onto an ozone-cleaned microscope coverslip, which was then dried in an oven at 50 ^o^C for 45 min. 25 μL of lipids (1 mg/mL) was subsequently spread onto the dried PVA film and placed under vacuum for 30 min until the solvent evaporated. 350 μL of sucrose solution (187 mM) was added to the PVA film and incubated for 45 min to allow GUV formation. The GUVs were then pipetted into an eppendorf tube and stored at 4 ^o^C until further use.

For coacervates-GUV interaction studies, 5 μL of the GUV solution was added to 200 μL of phosphate buffer and incubated for 5 min at room temperature. 20 μL of EGFP-loaded coacervates was then added to the GUVs for interaction studies. Fluorescence microscopy images were acquired using a Delta vision elite inverted epifluorescence microscope with Olympus IX-71 base fitted with 10×/0.40, 20×/0.75, 40x/0.65-1.35 oil objectives (Olympus, Tokyo, Japan), DAPI, TRITC and FITC Semrock filters (New York, NY), a mercury lamp (Intensilight C-HGFIE, Nikon Corporation, Tokyo, Japan), and a high-precision motorized stage. Images were collected using Softworx 4.1.0 (Applied Precision, Inc., Issaquah, WA) and processed using ImageJ.

#### Giant plasma membrane vesicles (GMPVs)

GPMVs were prepared using a previously described method by Gerstle et al., 2018. Briefly, HeLa cells that were grown to around 70% confluency in a T-25 cell culture flask, were gently rinsed twice with 1 mL vesiculation buffer (150 mM NaCl, 2 mM CaCl2, and 20 mM HEPES in water, pH 7.4) in order to remove dead cells and cell debris. 1 mL of the labelling buffer containing 1% (v/v) methanol, 2 μg/mL of Dil-C12 in vesiculation buffer was added to the flask and allowed to incubate at 37 ^o^C with occasional gentle rocking. At the end of the incubation, the labelling buffer was removed and the cells rinsed five times with vesiculation buffer. A final rinse was made using freshly prepared activation buffer (1.9 mM DTT and 27.6 mM formaldehyde in 10 mL of vesiculation buffer). 1 mL of activation buffer was then added to the cells and were allowed to incubate at 37 ^o^C for 90 min with occasional gentle rocking to separate GPMVs from cells. After 90 min, the flask was gently tapped from the bottom to release any cell-attached GPMVs and the solution containing the GPMVs were decanted into a 2 mL Eppendorf tube and allowed to sit for 30 min at RT. GPMVs for coacervate experiments were pipetted out from the bottom of the Eppendorf tube using a 1 mL pipette tip whose tip was cut with scissors to increase the size of the opening at the tip. For coacervates-GPMV interaction studies, 10 μL of the GPMV solution was added to 100 μL of EGFP-loaded coacervates previously pipetted on to a coverslip (glass bottom Mattek dishes). After gently pipetting the solution, the Mattek dishes were covered, sealed and the sample imaged immediately using a Carl Zeiss LSM 780 inverted microscope with a Plan-Apochromat 63x/1.4 oil objective. Images were collected using ZEN 2.3 SP1 FP3 (black) software as .czi files and were processed using FIJI.

#### Cell culture

HeLa and HepG2 cells (ATCC, USA) were cultured on DMEM (Gibco) and EMEM (ATCC) media supplemented with 10% FBS and penicillin/streptomycin solution (Gibco) in a humidified atmosphere at 37C° and 5% CO2. Cells were routinely tested for mycoplasma using Mycostrip kit (Invivogen).

#### Transfection of cells with coacervates for TEM and SEM

For cell transfection, reduced serum Optimem media (Gibco) was used. Coacervates were formed by mixing one part of peptide stock solution in 10 mM acetic acid (10 mg/ml for HB*pep*-SP or 20 mg/ml for HB*pep*) with 9 parts of 10 mM phosphate buffer with 100 mM sodium chloride, pH 7.5 for HB*pep* and 6.5 for HB*pep*-SP containing sfGFP (HB*pep*-SP peptide coacervates) or sfGFP and recombinant ferritin^39^ (HB*pep* coacervates). The media in cell culture dishes or flasks were replaced by Optimem, and coacervate mixtures were gently pipetted in. For HB*pep*, the final peptide concentration in Optimem during transfection was 0.4 mg/ml, for HB*pep*-SP - 0.1 mg/ml.

#### TEM

Cells were grown in Mattek dishes (Mattek) or T25 flasks (Corning) till ~60% confluence. For transfection, the media was substituted with Optimem (Gibco) containing coacervates loaded with EGFP (HB*pep*-SP peptide) or EGFP and ferritin (HB*pep*) as model cargos and placed in the incubator for 15 min or 3 h incubation followed by washing with PBS and fixation with 2.5% glutaraldehyde (EMS) in 0.1 M phosphate or cacodylate buffer, pH 7.4. Cells were postfixed with 1% OsO4 solution with added 2% potassium ferricyanide for 1.5 h on ice. Cells grown in Mattek dishes (HeLa transfected with HB*pep* coacervates) were embedded as monolayer while cells in T25 flasks. Other coacervates and cell lines were scraped after fixation and embedded as pellets. Cells in Mattek dishes were additionally stained with 1% of uranium acetate solution overnight in the dark. All samples were further subjected to dehydration through graded ethanol series and embedding in Durcupan resin. Ultrathin sections (70 or 100 nm) were obtained using Leica FC6 microtome. Imaging was performed on JEM-2100F electron microscope operated at 200kV in HAADF-STEM mode.

#### SEM

HeLa and HepG2 cells were cultured on glass coverslips in 6 well plates. Cells were transfected with EGFP-loaded coacervates followed by 15 min incubation at 37 °C in the incubator, washed with PBS, fixed with 4% paraformaldehyde in PBS for 30 min at room temperature and dehydrated in a graded ethanol series. Samples were CO2 critical point dried (Samdri-PVT-3D, Tousimis) and coated by sputtering with a 10 nm platinum layer. Imaging was performed on a field emission scanning electron microscope (SEI mode, 10-15keV, JSM-7600F, JEOL).

## Acknowledgements

We would like to acknowledge the Facility for Analysis, Characterisation, Testing and Simulation (FACTS), Nanyang Technological University (NTU), Singapore, for the use of their electron microscopy facilities and Dr YY. Tay for his advices and suggestions regarding TEM imaging. We thank the NTU Optical Bio-Imaging Centre (NOBIC) at the Singapore Centre for Environmental Life Sciences Engineering (SCELSE), NTU for the use of fluorescence and confocal microscopes. We also thank Prof. Atul Parikh (University of California, Davis) for helpful discussions. This research was funded by the Ministry of Education (MOE), Singapore, through an Academic Research Fund (AcRF) Tier 3 grant (Grant No. MOE 2019-T3-1-012).

## References

1. Sun, Y., Lim, Z. W., Guo, Q., Yu, J. & Miserez, A. Liquid–liquid phase separation of proteins and peptides derived from biological materials: Discovery, protein engineering, and emerging applications. MRS Bull. 45, 1039–1047 (2020).

2. Johnson, N. R. & Wang, Y. Coacervate delivery systems for proteins and small molecule drugs. Expert Opin. Drug Deliv. 11, 1829–1832 (2014).

3. Lim, Z. W., Ping, Y. & Miserez, A. Glucose-Responsive Peptide Coacervates with High Encapsulation Efficiency for Controlled Release of Insulin. Bioconjug. Chem. 29, 2176–2180 (2018).

4. Lim, Z. W., Varma, V. B., Ramanujan, R. V. & Miserez, A. Magnetically responsive peptide coacervates for dual hyperthermia and chemotherapy treatments of liver cancer. Acta Biomater. 110, 221–230 (2020).

5. Sun, Y. et al. Phase-separating peptides for direct cytosolic delivery and redox-activated release of macromolecular therapeutics. Nat. Chem. 14, 274–283 (2022).

6. Tan, Y. et al. Infiltration of chitin by protein coacervates defines the squid beak mechanical gradient. Nat. Chem. Biol. 11, 488–495 (2015).

7. Hou, X., Zaks, T., Langer, R. & Dong, Y. Lipid nanoparticles for mRNA delivery. Nat. Rev. Mater. 6, 1078–1094 (2021).

8. Behzadi, S. et al. Cellular uptake of nanoparticles: journey inside the cell. Chem. Soc. Rev. 46, 4218–4244 (2017).

9. Rennick, J. J., Johnston, A. P. R. & Parton, R. G. Key principles and methods for studying the endocytosis of biological and nanoparticle therapeutics. Nat. Nanotechnol. 16, 266–276 (2021).

10. Lim, J. P. & Gleeson, P. A. Macropinocytosis: an endocytic pathway for internalising large gulps. Immunol. Cell Biol. 89, 836–843 (2011).

11. Uribe-Querol, E. & Rosales, C. Phagocytosis: Our Current Understanding of a Universal Biological Process. Front. Immunol. 11, 1066 (2020).

12. Swanson, J. A. Shaping cups into phagosomes and macropinosomes. Nat. Rev. Mol. Cell Biol. 9, 639–649 (2008).

13. Mercer, J. & Helenius, A. Virus entry by macropinocytosis. Nat. Cell Biol. 11, 510–520 (2009).

14. Veiga, E. & Cossart, P. Listeria hijacks the clathrin-dependent endocytic machinery to invade mammalian cells. Nat. Cell Biol. 7, 894–900 (2005).

15. Ford, C., Nans, A., Boucrot, E. & Hayward, R. D. Chlamydia exploits filopodial capture and a macropinocytosis-like pathway for host cell entry. PLOS Pathog. 14, e1007051 (2018).

16. Iwata, T. et al. Liquid Droplet Formation and Facile Cytosolic Translocation of IgG in the Presence of Attenuated Cationic Amphiphilic Lytic Peptides. Angew. Chem. Int. Ed. 60, 19804–19812 (2021).

17. Gajendiran, M., Kim, S., Jo, H. & Kim, K. Fabrication of pH responsive coacervates using a polycation-b-polypropylene glycol diblock copolymer for versatile delivery platforms. J. Ind. Eng. Chem. 90, 36–46 (2020).

18. Chenglong, W. et al. Dextran-based coacervate nanodroplets as potential gene carriers for efficient cancer therapy. Carbohydr. Polym. 231, 115687 (2020).

19. Barthold, S. et al. Preparation of nanosized coacervates of positive and negative starch derivatives intended for pulmonary delivery of proteins. J. Mater. Chem. B 4, 2377–2386 (2016).

20. Armstrong, J. P. K. et al. Cell paintballing using optically targeted coacervate microdroplets. Chem. Sci. 6, 6106–6111 (2015).

21. Lu, T. et al. Endocytosis of Coacervates into Liposomes. J. Am. Chem. Soc. jacs.2c04096 (2022) doi:10.1021/jacs.2c04096.

22. Wimley, W. C. & White, S. H. Experimentally determined hydrophobicity scale for proteins at membrane interfaces. Nat. Struct. Mol. Biol. 3, 842–848 (1996).

23. Schillinger, A.-S., Grauffel, C., Khan, H. M., Halskau, Ø. & Reuter, N. Two homologous neutrophil serine proteases bind to POPC vesicles with different affinities: When aromatic amino acids matter. Biochim. Biophys. Acta BBA - Biomembr. 1838, 3191–3202 (2014).

24. de Jesus, A. J. & Allen, T. W. The role of tryptophan side chains in membrane protein anchoring and hydrophobic mismatch. Biochim. Biophys. Acta BBA - Biomembr. 1828, 864–876 (2013).

25. de Araujo, A. D., Hoang, H. N., Lim, J., Mak, J. Y. W. & Fairlie, D. P. Tuning Electrostatic and Hydrophobic Surfaces of Aromatic Rings to Enhance Membrane Association and Cell Uptake of Peptides. Angew. Chem. Int. Ed. 61, (2022).

26. Kruger, T. M. et al. Mechanosensitive Endocytosis of High-Stiffness, Submicron Microgels in Macrophage and Hepatocarcinoma Cell Lines. ACS Appl. Bio Mater. 1, 1254–1265 (2018).

27. Fenz, S. F. & Sengupta, K. Giant vesicles as cell models. Integr. Biol. 4, 982 (2012).

28. Andreev, K. et al. Hydrophobic interactions modulate antimicrobial peptoid selectivity towards anionic lipid membranes. Biochim. Biophys. Acta BBA - Biomembr. 1860, 1414–1423 (2018).

29. van Meer, G., Voelker, D. R. & Feigenson, G. W. Membrane lipids: where they are and how they behave. Nat. Rev. Mol. Cell Biol. 9, 112–124 (2008).

30. Sezgin, E. et al. Elucidating membrane structure and protein behavior using giant plasma membrane vesicles. Nat. Protoc. 7, 1042–1051 (2012).

31. Vercauteren, D. et al. The Use of Inhibitors to Study Endocytic Pathways of Gene Carriers: Optimization and Pitfalls. Mol. Ther. 18, 561–569 (2010).

32. Chaudhary, N. et al. Endocytic Crosstalk: Cavins, Caveolins, and Caveolae Regulate Clathrin-Independent Endocytosis. PLoS Biol. 12, e1001832 (2014).

33. Swanson, J. A. & Baer, S. C. Phagocytosis by zippers and triggers. Trends Cell Biol. 5, 89–93 (1995).

34. Pir Cakmak, F., Marianelli, A. M. & Keating, C. D. Phospholipid Membrane Formation Templated by Coacervate Droplets. Langmuir 37, 10366–10375 (2021).

35. Maity, A., De, S. K. & Chakraborty, A. Interaction of Aromatic Amino Acid-Functionalized Gold Nanoparticles with Lipid Bilayers: Insight into the Emergence of Novel Lipid Corona Formation. J. Phys. Chem. B 125, 2113–2123 (2021).

36. Yang, C. & Svitkina, T. Filopodia initiation: Focus on the Arp2/3 complex and formins. Cell Adhes. Migr. 5, 402–408 (2011).

37. Ingle, N. P., Hexum, J. K. & Reineke, T. M. Polyplexes Are Endocytosed by and Trafficked within Filopodia. Biomacromolecules 21, 1379–1392 (2020).

38. Mattila, P. K. & Lappalainen, P. Filopodia: molecular architecture and cellular functions. Nat. Rev. Mol. Cell Biol. 9, 446–454 (2008).

39. Sana, B. et al. The Role of Nonconserved Residues of Archaeoglobus fulgidus Ferritin on Its Unique Structure and Biophysical Properties. J. Biol. Chem. 288, 32663–32672 (2013).

