## Supplementary Materials for "Cellular Uptake of His-Rich Peptide Coacervates Occurs by a Macropinocytosis-Like Mechanism"

### Supplementary Information

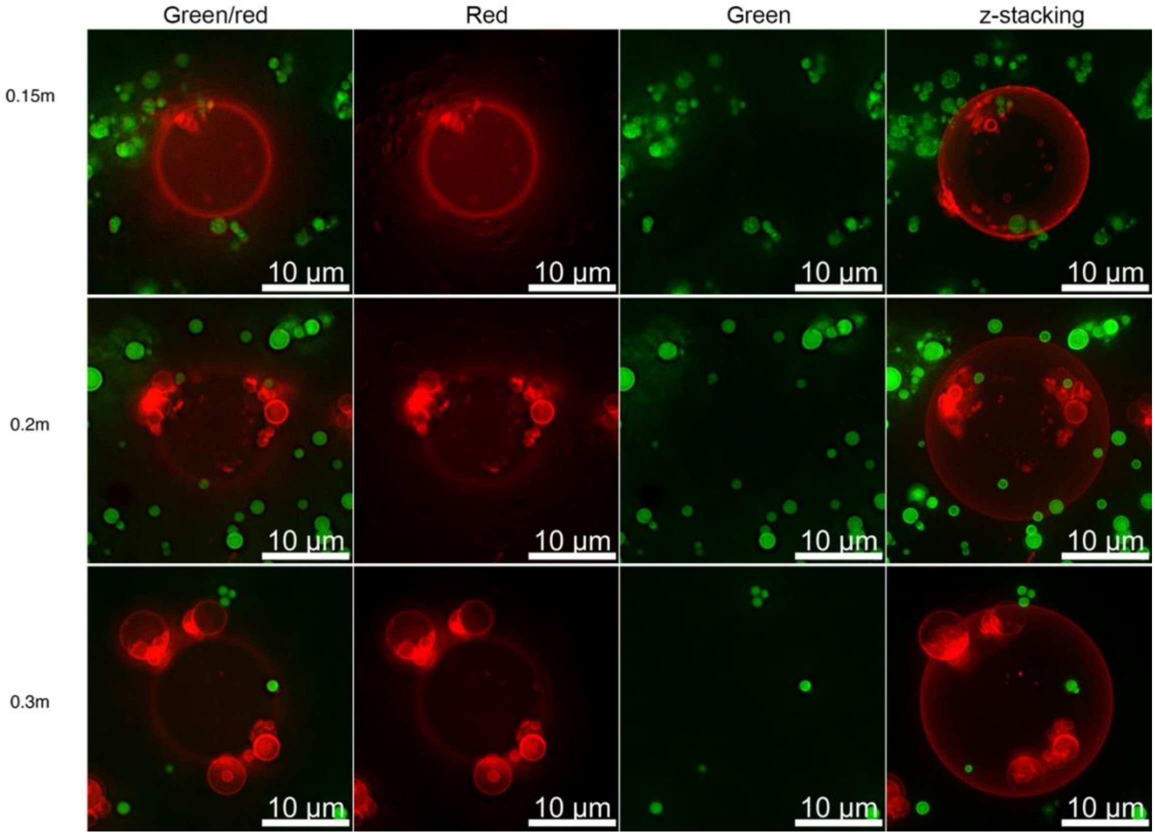


**Figure S 1. Effect of ionic strength on HB*pep* coacervate attachment to POPC GUVs.** Fluorescence microscopy images of POPC GUVs (Red) mixed with HB*pep* coacervates entrapping EGFP (green) at different ionic strengths. Representative merged images (left column) of the red and green channel (middle columns) are shown and the right-most column represents a representative z-stack composite of optical sections collected 1 µm apart.

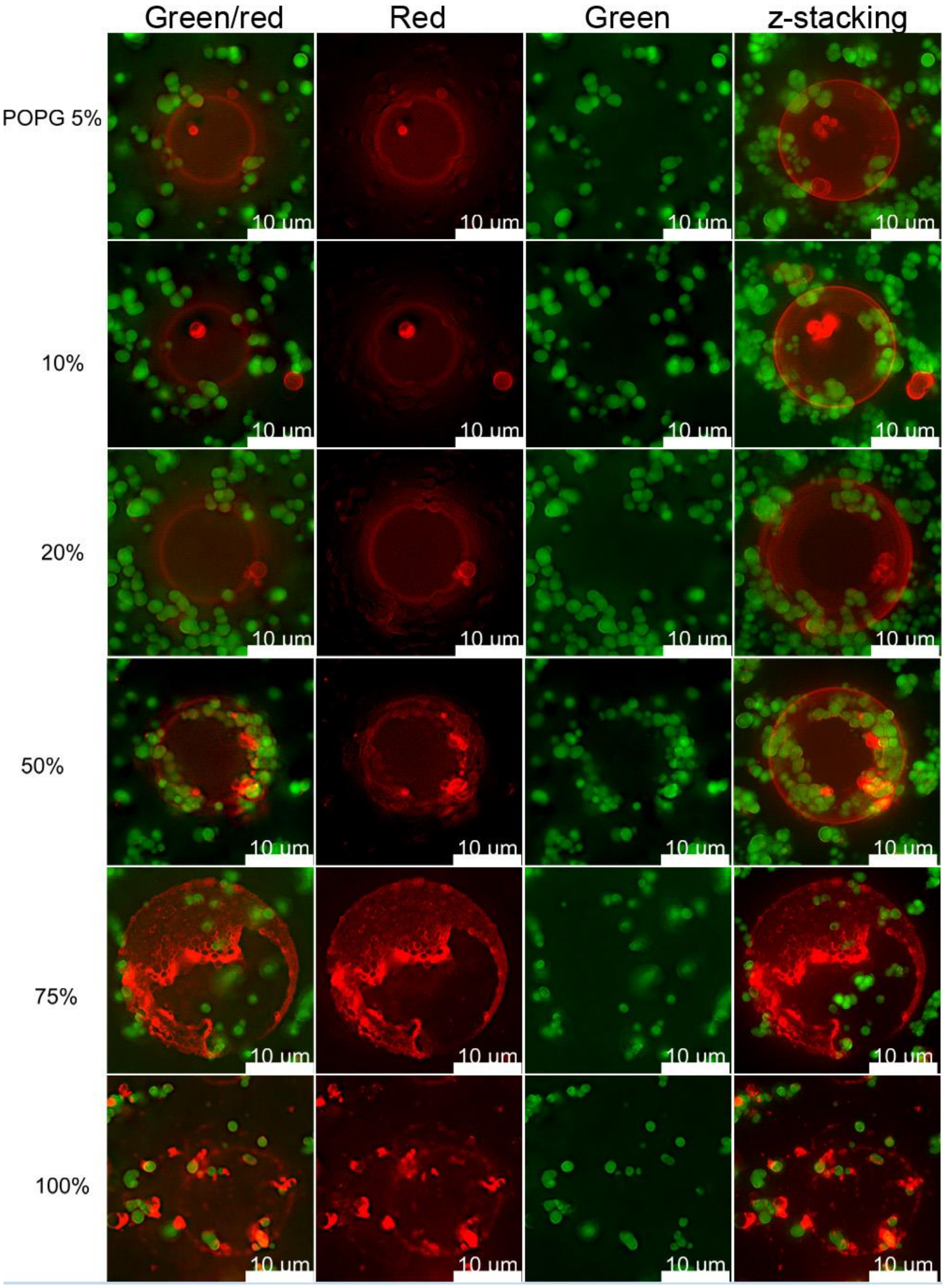


**Figure S 2. Effect of negatively charged membrane lipids on HB*pep* coacervate attachment.** Fluorescence microscopy images of POPC GUVs (Red) with varying levels of POPG is mixed with HB*pep* coacervates entrapping EGFP (green). Representative merged images (left column) of the green and red channel (middle columns) are shown and the right-most column represents a representative z-stack composite of optical sections collected 1 µm apart.

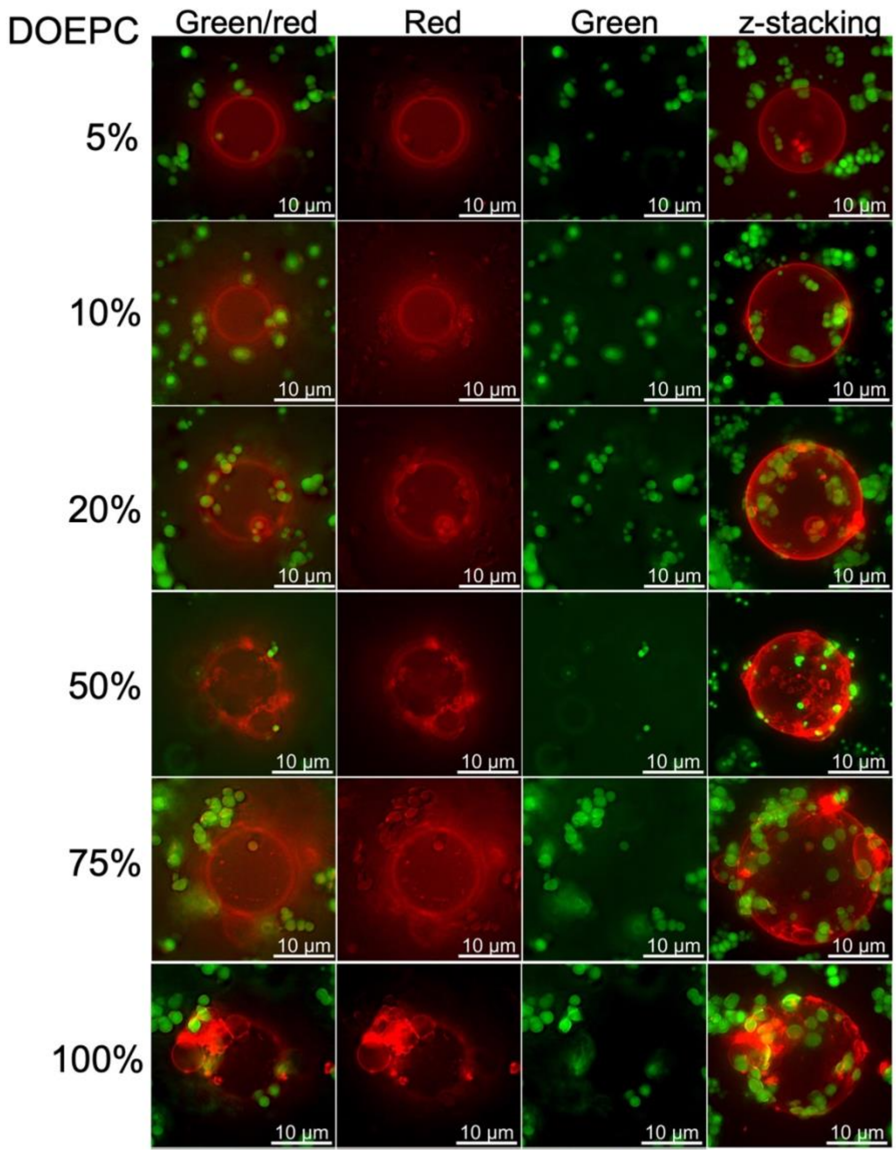


**Figure S 3. Effect of positively charged membrane lipids on HB*pep* coacervate attachment.** Fluorescence microscopy images of POPC GUVs (Red) with varying levels of DOEPC is mixed with HB*pep* coacervates entrapping EGFP (green). Representative merged images (left column) of the green and red channel (middle columns) are shown and the right-most column represents a representative z-stack composite of optical sections collected 1 µm apart.

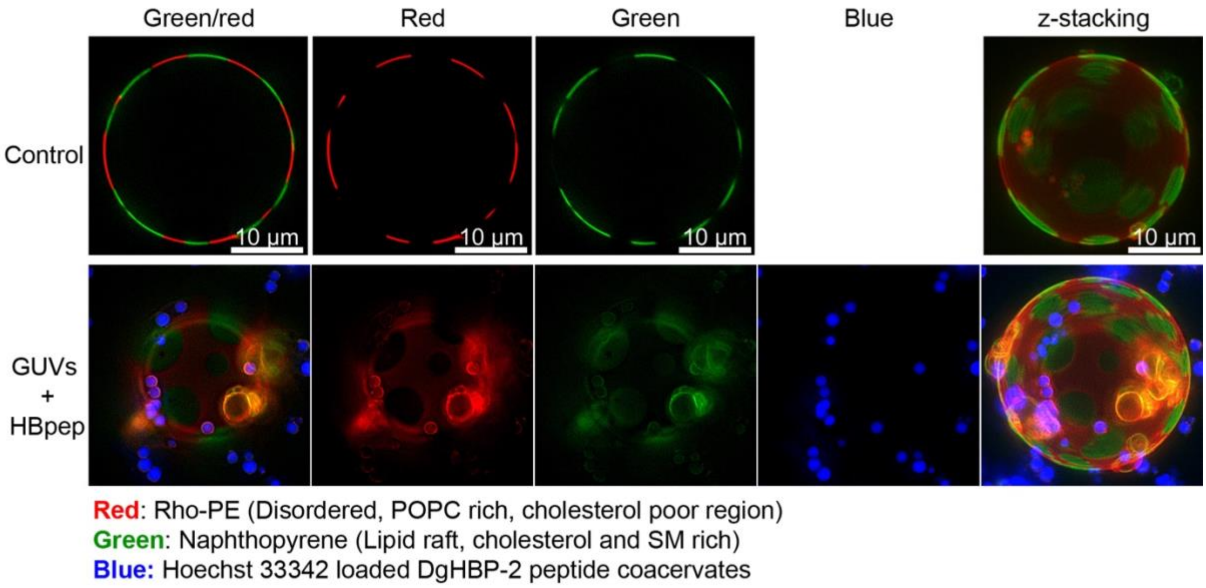


**Figure S 4. Interaction of HB*pep* coacervates with GUVs mimicking lipid rafts (phase separated liquid ordered and liquid disordered regions)**. Hoechst 33342-loaded HB*pep* coacervates mixed with GUVs prepared from 40% POPC/40% Sphingomyelin (SM)/20% Cholesterol in phosphate buffer. **Top row** (control): Disordered phase (Rho-PE, red) and ordered phase (Naphthopyrene, green) within a GUV. **Bottom row:** Attachment of Hoechst 33342-loaded coacervates with GUV containing lipid rafts.

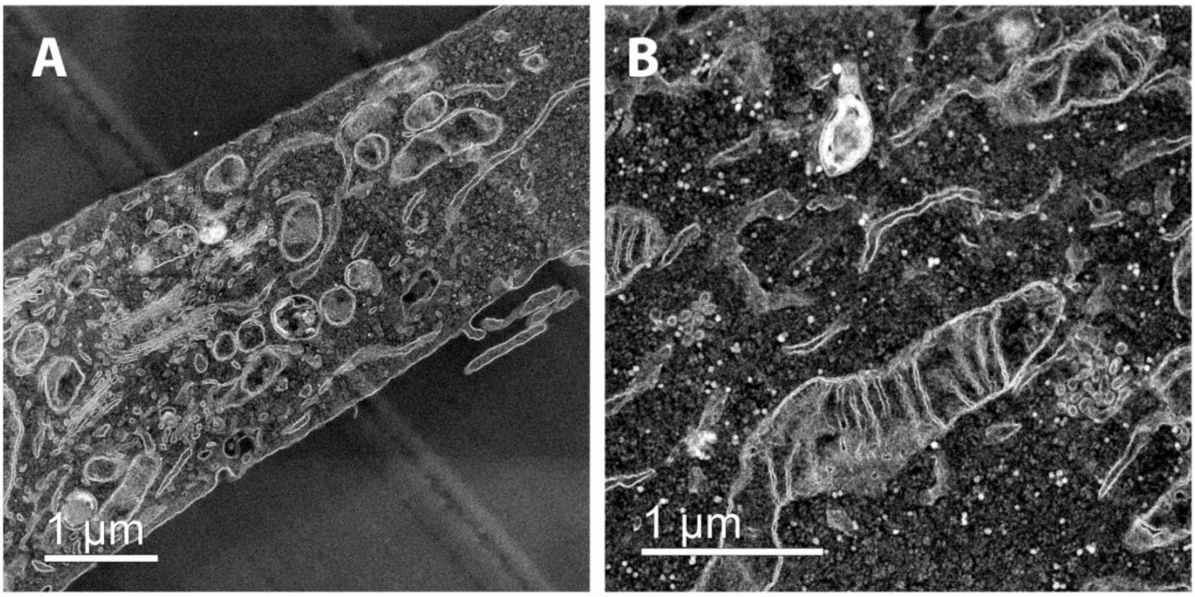


**Figure S 5. HAADF-STEM images of control HeLa cells not transfected with coacervates. (A)** A representative image of a cell without electron-dense spherical coacervates at the cell membrane and in the cytoplasm; **(B)** a close-up view of the cell cytoplasm demonstrating presence of electron dense lamellar body but not coacervates.

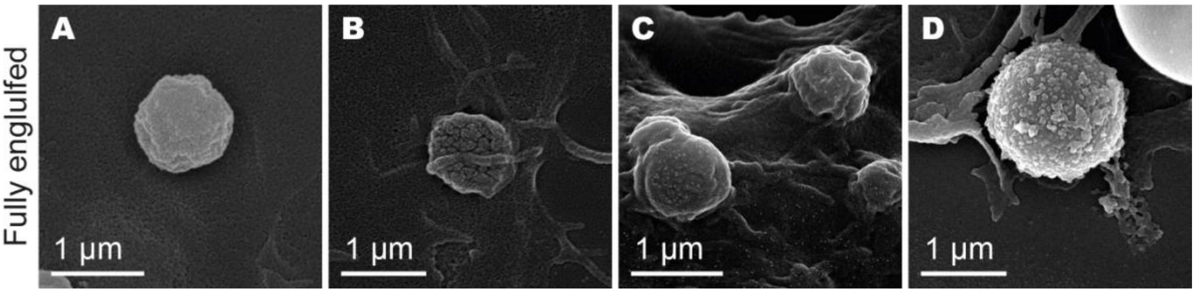


**Figure S 6. Representative SEM images of bumps found on the cell surface that may represent fully engulfed coacervates. (A)** HB*pep* coacervate in HeLa cell; **(B)** HB*pep*-SP coacervate in HeLa cell; **(C)** HB*pep* coacervate in HepG2 cell; **(D)** HB*pep*-SP coacervate in HepG2 cell.
